# Modernizing the original influenza vaccine to enhance protective antibody responses against neuraminidase antigens

**DOI:** 10.1101/2023.02.20.529183

**Authors:** Mira Rakic Martinez, Jin Gao, Hongquan Wan, Hyeog Kang, Laura Klenow, Robert Daniels

## Abstract

Most seasonal influenza vaccines are produced using hemagglutinin (HA) surface antigens from inactivated virions. However, virions are thought to be a suboptimal source for the less abundant NA surface antigen, which is also protective against severe disease. Here, we demonstrate that inactivated influenza virions are compatible with two modern approaches for improving protective NA antibody responses. Using a DBA/2J mouse model, we confirmed that the strong infection-induced NA inhibitory (NAI) antibody responses are only achieved by high dose immunizations of inactivated virions, likely due to low viral NA content. Based on this observation, we first produced virions with higher NA content by using reverse genetics to exchange the viral internal gene segments. Single immunizations with these inactivated virions enhanced NAI antibody responses, improved NA-based protection from a lethal viral challenge and allowed the development of natural immunity to the heterotypic challenge virus HA. Second, we combined inactivated virions with recombinant NA proteins. These mixtures increased NA-based protection following viral challenge and elicited stronger NA antibody responses than either component alone, especially when the NAs were homologous. Together, these results indicate that viral- and protein-based vaccines can be combined in a single syringe to improve protective antibody responses to influenza antigens.

## INTRODUCTION

Influenza vaccines were introduced shortly after methods for propagating and purifying the virus were established using embryonated eggs in the 1930s. The initial vaccine was comprised of formalin inactivated virions from a single H1N1 strain (A/PR/8/34) of influenza A virus (IAV) [1]. Following discovery of influenza B viruses (IBVs), it was updated to a bivalent vaccine consisting of inactivated virions from both an IAV and IBV strain [2]. Decades later, inactivated virions were treated with detergent (or ether) to create split vaccines that elicit lower reactogenicity following administration [3] and are still commonly used today. More recently, influenza vaccines have been introduced that use viruses propagated in mammalian cells [4], recombinant hemagglutinin (HA) antigens produced by insect cells [5], or infectious attenuated viruses produced in eggs [6]. Independent of the platform, all influenza vaccines are currently available as quadrivalent formulations that are frequently updated to include HA antigens from recently circulating IAV (H1N1 and H3N2) and IBV (Yamagata and Victoria lineages) strains [7]. Despite these improvements, seasonal influenza vaccine efficacy against disease remains suboptimal and season dependent.

All currently approved influenza vaccines were primarily developed to elicit optimal antibody responses against the HA antigens in the recommended influenza A and B vaccine strains. However, strain recommendations are made months in advance and influenza viruses are constantly evolving resulting in changes to HA that can periodically cause antigenic drift [8]. Consequently, HAs in circulating strains can become antigenically distinct from those in the recommended strains resulting in decreased seasonal vaccine efficacy [9]. One approach for improving influenza vaccine efficacy is to alleviate this HA dependency by including additional antigens such as neuraminidase (NA), which has been shown to reduce severe disease in animal models and humans challenged with infectious virus [10-14].

The primary role of NA is to promote viral movement through the respiratory mucosa by removing local sialic acid ligands that facilitate binding by the HA molecules on the viral surface [15]. Not surprisingly, antibodies that inhibit the ability of NA to cleave sialic acid have been shown to correlate with protection [16, 17] and reduce viral replication and shedding [18], similar to chemical NA inhibitors such as oseltamivir [19, 20]. NA protective antibodies have also been isolated from humans that do not inhibit its activity [14], indicating antibodies against NA can provide protection by two or more mechanisms. In contrast to HA, NA only has nine subtypes (N1-N9) in the avian IAV reservoir, and its antigenic drift appears to be discordant from HA [21], suggesting a vaccine against HA and NA would provide broader strain coverage, potentially resulting in higher efficacy.

While viral-based vaccines possess NA, influenza viruses by nature incorporate low NA amounts and only a few studies have assessed NA antigenicity in circulating strains [14, 21-23]. In addition, NA quantity and quality in viral-based vaccines is not regulated and a recent study has shown that split vaccine responses are heavily biased towards HA [24]. Seminal work demonstrated that NA is a protective antigen in influenza virus-challenged humans that either possessed NA antibodies from prior infection [11] or were immunized with inactivated IAV virions that contained an identical NA, but a different HA subtype [12]. Later studies in animals used NA isolated from viruses to suggest influenza vaccines can be combined with additional NA antigen to increase the NA protective response [25]. However, supplementation of influenza vaccines with NA has not been realized almost 25 years later as most studies have independently focused on NA-based protection achieved by immunization with recombinant NA (rNA) antigens [26-33]. More recent work has shown that NA can potentially be combined with HA and other influenza antigens in mRNA vaccines as well [34, 35], but these vaccines are still in development. Despite the numerous types of influenza vaccines, viruses remain the most common antigen source. However, viruses are also presumed to be nonideal for NA antigens due to the low content. Here, we re-evaluated inactivated virions as a possible platform for eliciting protective NA antibody responses because they present NA in a large multivalent format and eliminate any potential NA loss due to split vaccine manufacturing that was widely implemented during the time of the seminal work on NA. Our results show that although high doses of inactivated virions are required to elicit the strong NA antibody responses that are observed following viral infection, NA-based protection in mice was achieved with single doses of inactivated virions that were much lower than those required for a recombinant NA antigen. We lowered the protective dose further (∼4-fold) by exchanging the viral internal genes to increase NA content and confirmed that the NA-based protection decreased severity while enabling natural immunity to develop against the challenge virus. Lastly, we demonstrated inactivated virions can be combined with a homotypic or heterotypic rNA protein in a single vaccine dose to enhance NA protective responses more than either component individually. Together, these data indicate that inactivated virions offer more flexibility for eliciting protective NA responses than previously acknowledged and suggest that virions are amenable to supplementation with rNA antigens. The potential benefits of combining inactivated virions with recombinant influenza antigens is discussed.

## RESULTS

### Only high doses of inactivated virions elicit strong infection-like NA antibody responses

Previous studies indicate influenza virus infections elicit stronger and more frequent NA antibody responses than influenza vaccines either because the NA amount in the vaccine is insufficient, or the native structure is disrupted [24, 36]. To examine these possibilities, we compared NA and HA antibody responses in mice following infection and after immunization with β-propiolactone (BPL) inactivated virions (Fig. 1A), which were chosen to limit potential NA denaturation. Initially, groups of female DBA/2J mice were infected with ten-fold dilutions of a BR18^H1N1^ virus or a recombinant PR8^H6N1-BR18^ virus that both carry identical NA antigens (N1-BR18) and different HA antigens (subtypes H1 and H6). Sera from mice that received the highest virus amount without exhibiting 25% weight loss were obtained 2 weeks post-infection and analyzed (Fig. 1B). NA inhibitory (NAI) titres (Fig. 1C, left panel), determined by an enzyme-linked lectin assay [23], and HA agglutination inhibition (HAI) titres (Fig. 1C, right panel) were both higher in the mice that recovered from the BR18^H1N1^ infection opposed to the PR8^H6N1-BR18^ infection. Despite the differences, NAI to HAI titre ratios ranged from 10 to 20 independent of the viruses used for the infection or assays (Fig. 1D), indicating mice develop high NAI titres at two weeks post-infection with these viruses.

**Figure 1.**
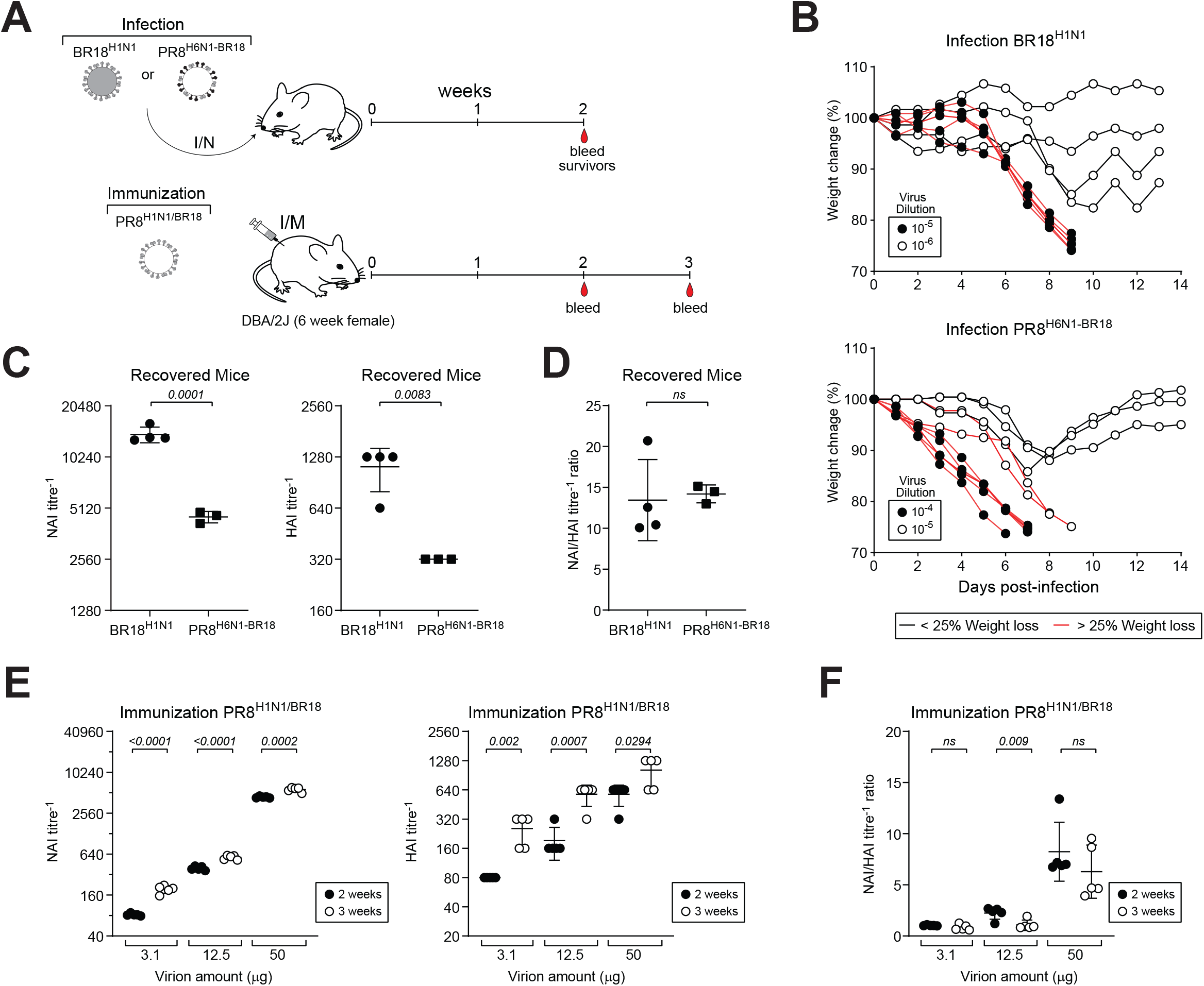
Comparison of NA antibody responses from infection and immunization with inactivated virions. **A**. Diagram of the approach used to compare NA antibody responses in mice following intranasal (I/N) infection with BR18^H1N1^ or PR8^H6N1-BR18^ viruses and intramuscular (I/M) immunization with inactivated PR8^H1N1/BR18^ virions. **B**. Graphs displaying weight loss in the individual mice following infection with the indicated BR18^H1N1^ (top panel) and PR8^H6N1-BR18^ (bottom panel) virus dilutions. Mice that lost ∼25% of initial body weight were euthanized. **C**. NAI (left panel) and HAI (right panel) titres are shown for the mice that survived the BR18^H1N1^ (4 mice) or PR8^H6N1-BR18^ (3 mice) infections. NAI titres from BR18^H1N1^ and PR8^H6N1-BR18^ infections were determined with PR8^H6N1-BR18^ and BR18^H1N1^ viruses, respectively. HAI titres were determined with the same virus used for infection. **D**. Graph displaying the NAI/HAI titre ratios in each mouse. **E**. NAI (left panel) and HAI (right panel) titres are shown for each mouse at 2- and 3-weeks post-immunization with the indicated amounts of inactivated PR8^H1N1/BR18^ virions. PR8^H6N1-BR18^ and PR8^H1N1/BR18^ viruses were used to determine NAI and HAI titres, respectively. **F**. NAI/HAI titre ratios are shown for the individual mice at 2- and 3-weeks post-immunization. *P* values (95% CI) are from a two-tailed unpaired t-test.

Next, we immunized mice with three different doses of inactivated PR8 ^H1N1/BR18^ virions that carry the same NA and HA antigens as BR18^H1N1^ (Fig. 1A). All mice displayed dose-dependent NAI and HAI antibody responses at both 2- and 3-weeks post-immunization and the titres were consistently higher at 3 weeks (Fig. 1E). At both times, only the high dose of BPL inactivated virions (50 μg) elicited NAI titres and NAI to HAI titre ratios that were observed following recovery from infection (Fig. 1F). As current vaccines are standardized to HA content (∼15 μg per HA per human dose), these data suggest that even with optimal conditions for stabilizing the NA structure, the low NA content limits the ability of viral-based vaccines to elicit NA antibody responses that are similar to natural infection.

### Manipulating virions to increase NA content reduces the dose needed for NA-based protection

Influenza candidate vaccine viruses (CVVs) are often high growth PR8 reassortants that are selected based on HA content [37]. Recently, we showed that virion NA content can be increased by exchanging the PR8 (A/PR/08/34) backbone or polymerase genes with those from WSN (A/WSN/1933), another high growth lab strain [38]. However, we did not test if the achieved NA content increase effectively reduced the inactivated virion amount that is needed in a single dose for NA-based protection from a lethal viral challenge. To address this question, we isolated and inactivated WSN^H1N1/BR18^ and PR8^H1N1/BR18^ virions that both contain the NA and HA antigens from BR18^H1N1^, which are identical to the recommended H1N1 vaccine antigens for the 2019-2020 influenza season (Fig.2A). Analysis of equal total protein amounts by Coomassie stained gel and NA activity (Fig. 2A) indicated that the WSN^H1N1/BR18^ virion preparation contained ∼4-fold more NA protein than the PR8^H1N1/BR18^ virion preparation with a slight reduction in HA content, in line with our previous study [38]. Mice were then immunized with different amounts of the inactivated virions and serological analysis was performed at 3 weeks, just prior to a lethal challenge with a PR8^H6N1-BR18^ reassortant virus that possesses the same NA (N1-BR18) as BR18^H1N1^ and a different HA (H6) subtype (Fig. 2B).

**Figure 2.**
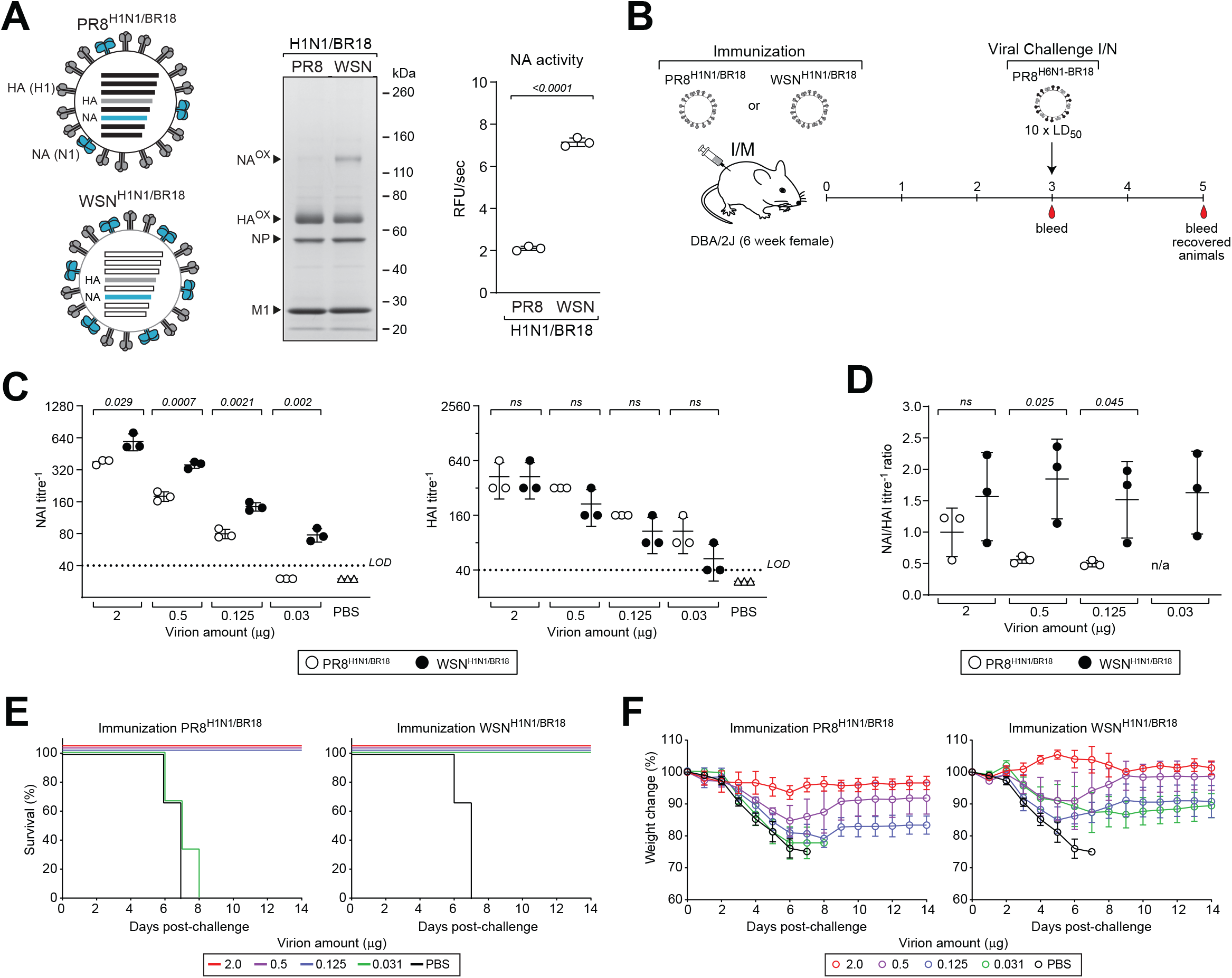
Inactivated virion NA content determines the dose needed for NA-based protection. **A**. Schematic of PR8^H1N1/BR18^ and WSN^H1N1/BR18^ virions (left) that contain different amounts of the same NA and HA antigens, non-reducing Coomassie stained gel (middle panel) where 4 μg of the BPL inactivated virions were resolved, relative NA activities (right panel) in 100 ng of BPL inactivated virions. **B**. Diagram of the immunization and challenge approach for comparing NA antibody responses and protection provided by the BPL inactivated PR8^H1N1/BR18^ and WSN^H1N1/BR18^ virions. **C**. NAI (left panel) and HAI (right panel) titres are shown for each group of mice (n=3) at 3-weeks post-immunization with the indicated amounts of inactivated PR8^H1N1/BR18^ and WSN^H1N1/BR18^ virions. PR8^H6N1-BR18^ and PR8^H1N1/BR18^ viruses were used to determine NAI and HAI titres, respectively. **D**. NAI/HAI titre ratios are shown for the individual mice in each group 3-weeks post-immunization. **E**. Graph displaying the percentage of mice from each vaccine group that survived the PR8^H6N1-BR18^ virus challenge. **F**. Body weight was monitored in all the mice following viral challenge and the mean for each vaccine group is displayed ± s.d. *P* values (95% CI) are from a two-tailed unpaired t-test.

At each dose the inactivated WSN^H1N1/BR18^ virions elicited higher NAI titres than the PR8^H1N1/BR18^ virions (Fig. 2C, left panel) and the HAI titres were equal or slightly lower (Fig. 2C, right panel), resulting in higher NAI/HAI titre ratios (Fig. 2D). Following viral challenge, protection was observed from all four doses (0.03 μg, 0.125 μg, 0.5 μg, and 2 μg) of the inactivated WSN^H1N1/BR18^ virions, but only the three higher doses of the PR8^H1N1/BR18^ virions (Fig. 2E), suggesting the difference is related to NA viral content. Mice that received the inactivated WSN^H1N1/BR18^ virions also showed less weight loss and faster recovery to a higher initial weight than those that received equivalent amounts of inactivated PR8^H1N1/BR18^ virions (Fig. 2F). These results, collected from the same experiment, demonstrate that NA virion content can be manipulated to significantly decrease (∼4-fold) the amount of inactivated virions that is required for NA-based protection in mice from a single dose.

### Virion-based NA protection enables natural immunity to heterotypic challenge virus HAs

The lack of sterilizing immunity from NA antigens has previously been shown to enable the development of natural immunity against infecting viral strains [39]. As this property can be advantageous when HAs in vaccine and circulating strains are antigenically mismatched, we examined the NAI and HAI responses in all immunized mice that recovered from the PR8^H6N1-BR18^ viral challenge at 2 weeks post-challenge, 5 weeks after immunization (Fig. 3A). Independent of the inactivated virions used for immunization all mice showed higher NAI titres against N1-BR18 following recovery and the increase retained the dose-dependent trend that was observed following immunization and pre-challenge. All mice also exhibited relatively unchanged HAI titres against the H1 antigen in the virions following recovery (Fig. 3B), but higher HAI titres against the challenge virus H6 antigen (Fig. 3C) which differed between virions. In mice immunized with the PR8^H1N1/BR18^ virions all the HAI titres against H6 significantly increased to similar levels following recovery, whereas mice that received the higher WSN^H1N1/BR18^ virion doses displayed decreased HAI (H6) titres, likely due to the higher NA antibody responses limiting the infection severity (Fig. 2). Despite the lower HAI (H6) titre, the response was still significant in mice that received the 2 μg dose of WSN^H1N1/BR18^ virions and displayed no weight loss following challenge. These results support the notion that NA-based protection from single doses of inactivated virions can limit infection severity and allow for the development of natural immunity to a heterotypic HA (antigenically mismatched) from a lethal influenza virus.

**Figure 3.**
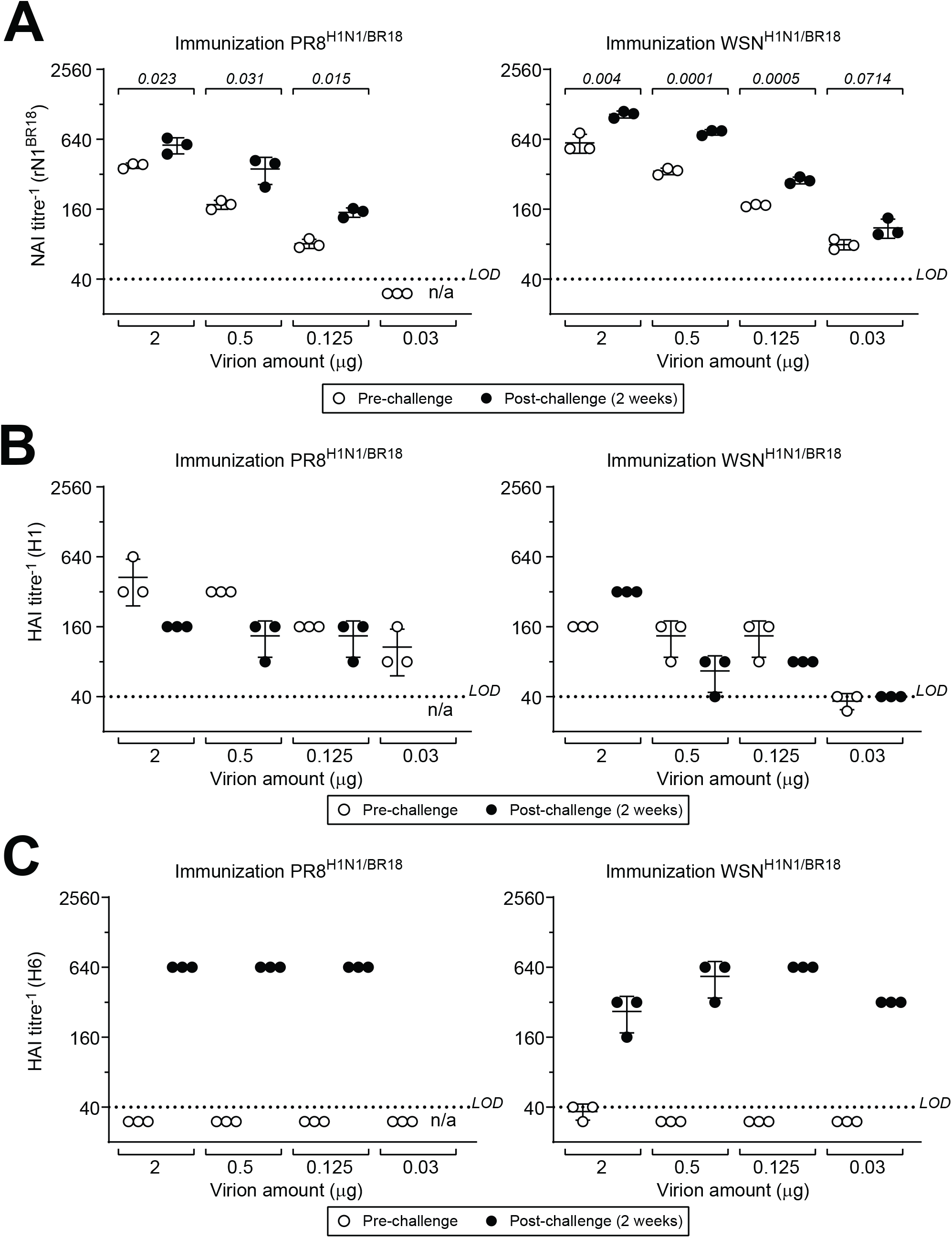
NA-based protection enables natural immunity to the challenge virus HA. **A**. NAI titres are shown for each group of vaccinated mice (n=3) pre-challenge (3-weeks post-immunization with inactivated virions) and 2-weeks post-challenge with the PR8^H6N1-BR18^ virus. Titres were determined using rN1^BR18^ protein to prevent interference from HA antibodies. Note all mice immunized with 0.03 μg of PR8^H1N1/BR18^ virions succumbed to the challenge. n/a - not applicable. **B**. HAI titres against H1 are shown for the same groups of mice pre-challenge and 2-weeks post-challenge with the PR8^H6N1-BR18^ virus. Titres were determined using the BR18^H1N1^ virus. **C**. HAI titres against the H6 antigen in the challenge virus are shown for the same groups of mice pre-challenge and 2-weeks post-challenge with the PR8^H6N1-BR18^ virus. Titres were determined using the PR8^H6N1-BR18^ virus.

### Mixing rNAs with inactivated virions improves NA antibody responses and protection

Increasing the NA virion content by exchanging the internal gene segments caused a corresponding decrease in HA virion content that can potentially reduce the number of vaccine doses. Therefore, we asked if protective NA antibody responses could be improved by mixing BPL inactivated PR8^H1N1/BR18^ virions with rNA protein (rN1^BR18^) from the same H1N1 strain (Fig. 4A). For this analysis we chose a dose (0.03 μg) of inactivated PR8^H1N1/BR18^ virions which protected against a lethal virus challenge with BR18^H1N1^ that carries the same NA and HA antigens (Supplemental Fig. 1), but not PR8^H6N1-BR18^ which only carries the same NA antigen (Figs. 2E and F).

**Figure 4.**
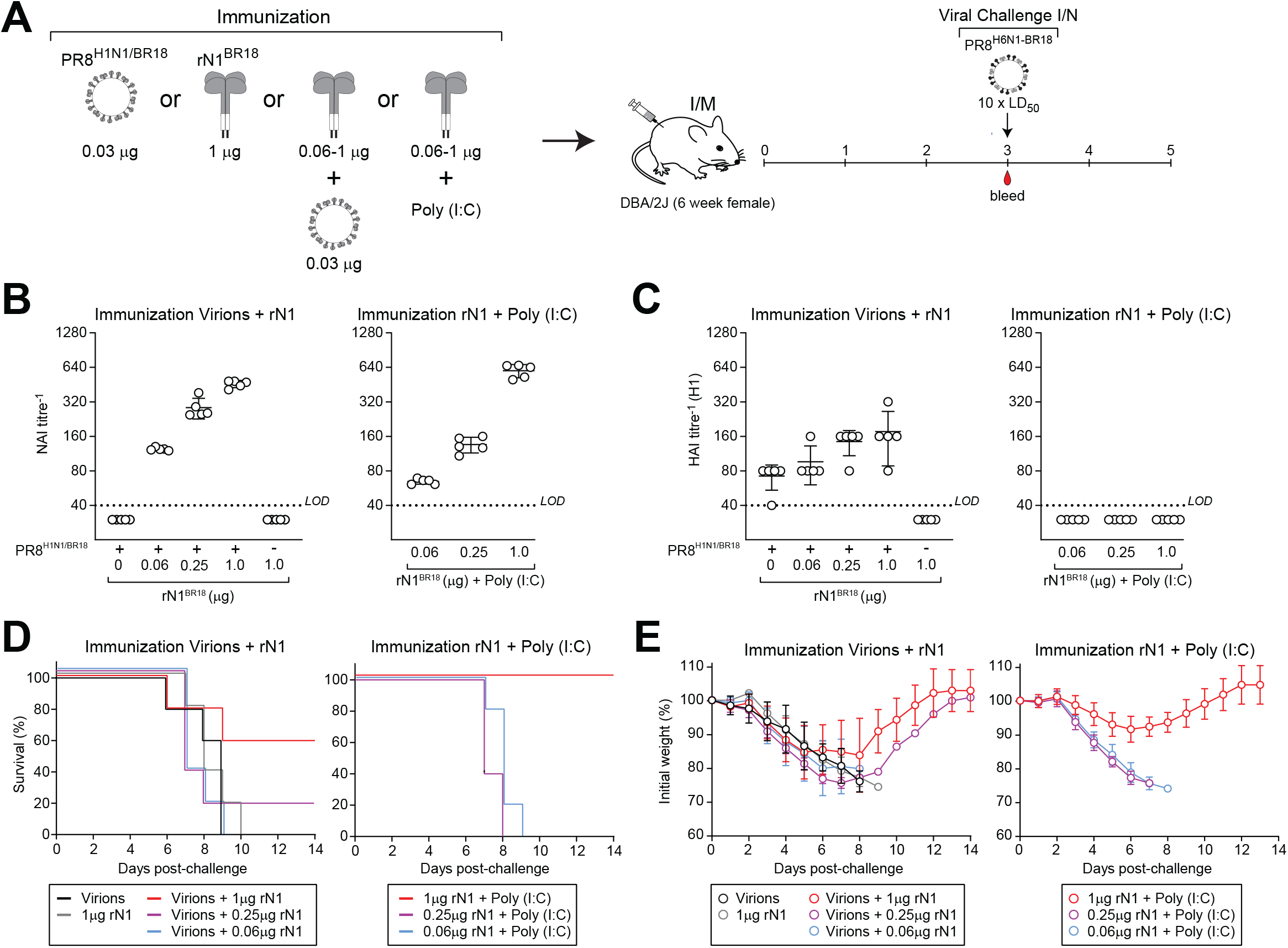
Mixing inactivated virions with rNA enhances protective NA antibody responses. **A**. Diagram of the immunization groups and challenge approach for examining in NA antibody responses are enhanced by mixing low amounts of BPL inactivated PR8^H1N1/BR18^ virions with rN1^BR18^ protein. **B**. NAI titres are shown for each group of mice (n=5) 3-weeks post-immunization with 0.03 μg of BPL inactivated PR8^H1N1/BR18^ virions mixed with the indicated rN1^BR18^ protein amounts, rN1^BR18^ protein alone, or the indicated rN1^BR18^ protein amounts mixed with the adjuvant poly (I:C). Titres were determined with the PR8^H6N1-BR18^ virus. **C**. HAI titres against H1 are shown for the same groups of mice 3-weeks post-immunization. Titres were determined using the BR18^H1N1^ virus. **D**. Graph displaying the percentage of mice from the indicated vaccine groups that survived the PR8^H6N1-BR18^ virus challenge. **E**. Body weight was monitored in all mice following viral challenge and the mean for each vaccine group is displayed ± s.d. *P* values (95% CI) are from a two-tailed unpaired t-test.

A characterization analysis confirmed the rN1^BR18^ protein is predominantly tetrameric and enzymatically active with a full-length purity of ∼85% (Supplemental Fig. 2). Based on previous work in our naïve mouse model with rN1^BR18^ proteins [27], we immunized groups of mice with the inactivated virions mixed with three different amounts of rN1^BR18^ protein (1 μg, 0.25 μg, or 0.0625 μg) and challenged them 3 weeks later with the PR8^H6N1-BR18^ virus (Fig. 4A). Inactivated virions alone and rN1^BR18^ protein with and without poly (I:C) adjuvant were included as controls.

All control groups displayed NAI titres below the limit of detection, whereas the groups mice that received inactivated virions mixed with rN1^BR18^ protein or poly (I:C) adjuvanted rN1^BR18^ protein showed dose-dependent NAI titres (Fig. 4B). Groups that received the virion and rN1^BR18^ protein combination also exhibited unexpected elevations in HAI (H1) titres compared to the virions alone (Fig. 4C), suggesting that the enzymatic activity, rN1^BR18^ protein, or other substances in the preparation stimulated the response to the HA antigen in the inactivated virions.

Following the PR8^H6N1-BR18^ virus challenge some of the mice that received inactivated virions mixed with either 0.25 or 1 μg of rN1^BR18^ protein survived, whereas all mice from the control groups succumbed to the challenge, with the exception of the high dose (1 μg) rN1^BR18^ poly (I:C) adjuvanted group (Fig. 4D). This finding is in line with our previous work [27] and this group also showed less weight loss than the group that received the combination of inactivated virions and 1 μg of rN1^BR18^ (Fig. 4E). While these data indicate the NA-based protection would have benefited from adding higher rN1^BR18^ protein amounts to the virions, they also suggest that the inactivated PR8^H1N1/BR18^ virions and the rN1^BR18^ both contribute to the NA antigen content.

To investigated if inactivate virions can also stimulate rNA responses, mice were immunized with PR8^H1N1/BR18^ virions mixed with 1 or 4 μg of either the homologous rN1^BR18^ or a heterologous rNA (rN2^KS17^) and challenged with a virus that carries a NA identical to the rNA (Fig. 5A). As previously seen, combining inactivated virions with rN1^BR18^ induced elevated NAI and HAI (H1) titres (Fig. 5B), which correlated with high survival rates (80%) following the PR8^H6N1-BR18^ challenge and a dose-dependent recovery in weight loss (Fig. 5C). Protection was only observed in the control group that received 4 μg rN1^BR18^ and it was higher than the group that received inactivated virions combined with 4 μg of rN1^BR18^ (Fig. 5C, left panel). However, the weight loss in this control group was more pronounced (Fig. 5C, right panel).

**Figure 5.**
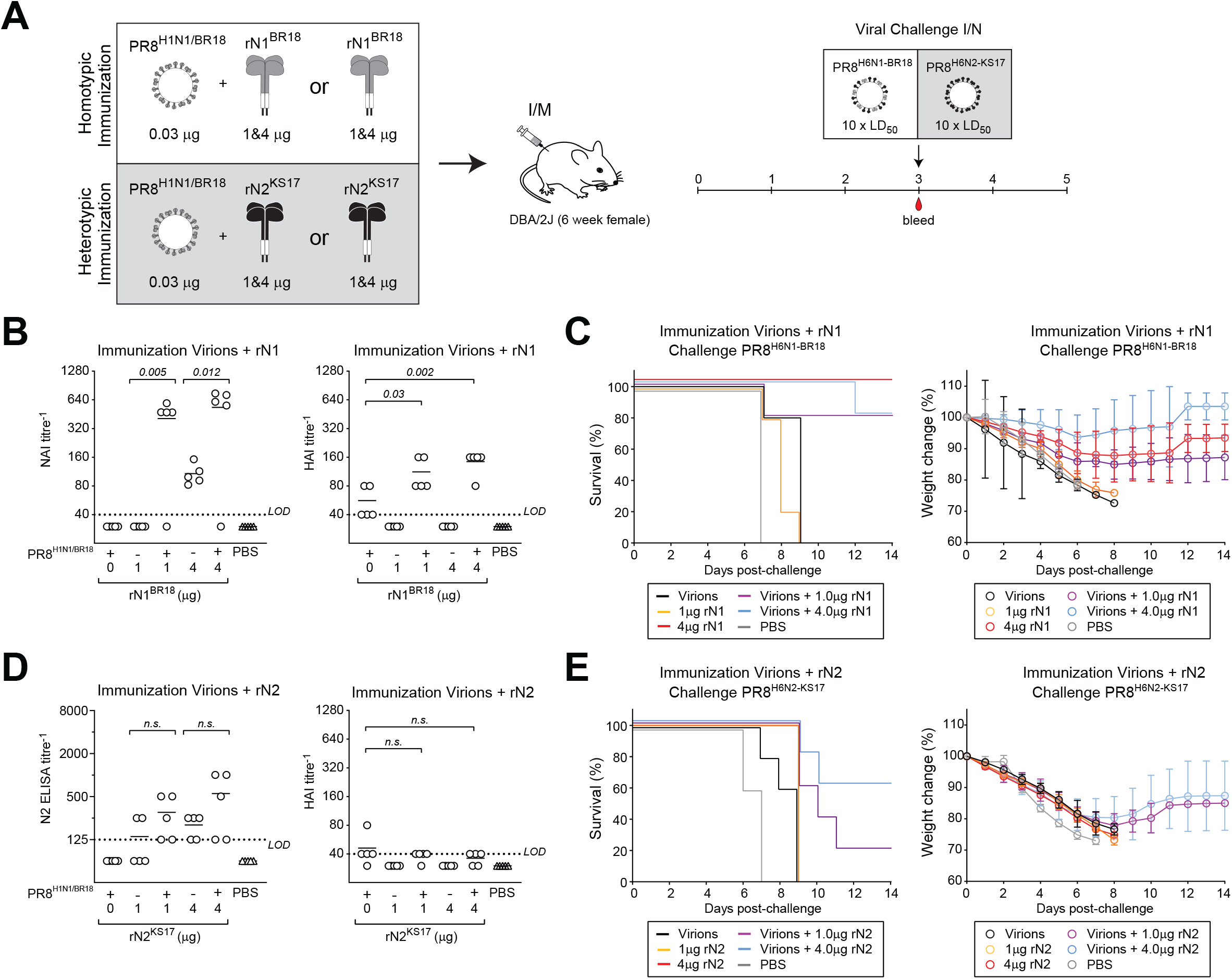
Immune responses and protection from inactivated virions mixed with homologous or heterologous rNAs. **A**. Immunization and challenge scheme for comparing antibody and protective responses against a homotypic (rN1^BR18^) and a heterotypic (rN2^KS17^) rNA mixed with a low amount of BPL inactivated PR8^H1N1/BR18^ virions. **B**. Graphs showing NAI (left panel) and HAI (right panel) titres obtained from each group of vaccinated mice (n=5) 3-weeks post-immunization with 0.03 μg of BPL inactivated PR8^H1N1/BR18^ virions mixed with the indicated amounts of rN1^BR18^ protein, or rN1^BR18^ protein alone. NAI and HAI titres were determined with PR8^H6N1-BR18^ and PR8^H1N1/BR18^ viruses, respectively. **C**. The percentage of mice from each vaccinated group that survived the PR8^H6N1-BR18^ virus challenge (left panel) is displayed with the mean body weight of each group ± s.d (right panel). **D**. Graphs showing NAI (left panel) and HAI (right panel) titres obtained from each group of vaccinated mice (n=5) 3-weeks post-immunization with 0.03 μg of BPL inactivated PR8^H1N1/BR18^ virions mixed with the indicated rN2^KS17^ protein amounts, or rN2^KS17^ protein alone. NAI and HAI titres were determined with PR8^H6N2-KS17^ and PR8^H1N1/BR18^ viruses, respectively. **E**. The percentage of mice from each vaccinated group that survived the PR8^H6N2-KS17^ virus challenge (left panel) is displayed with the mean body weight of each group ± s.d (right panel). *P* values (95% CI) are from a two-tailed unpaired t-test.

Similar but more subtle trends were observed with the inactivated PR8^H1N1/BR18^ virions and the rN2^KS17^ protein, which was shown to possess similar quality attributes to the rN1^BR18^ protein (Supplemental Fig. 2). In both groups that received the mixture of inactivated PR8^H1N1/BR18^ virions and rN2^KS17^, some mice exhibited higher N2 antibody titres than the control groups (Fig. 5D, left panel), but no increase was observed in HAI (H1) titres (Fig. 5D, right panel). Following the PR8^H6N2-KS17^ virus challenge, only groups that received the mixture showed some protection (Fig. 5E) and the surviving mice possessed the highest N2 antibody titres. Together, these results demonstrate that inactivated virions combined with a homotypic or heterotypic rNA are more protective than either component alone and that inactivated virions likely have some immunostimulatory effect on rNA antigens.

## DISCUSSION

Although human trials have shown NA to be a protective and potentially safe antigen [11, 12, 40], vaccines have still not been developed that effectively elicit NA antibody responses. Most studies on viral-based vaccines have attributed poor NA responses to the viral content being lower than HA, immunodominance, and variable loss of the more labile NA during manufacturing [26, 41-44]. However, these studies were mainly performed on split vaccines that were widely adopted around the time of the first NA human trials. Here, we hypothesized the original inactivated virion vaccines may effectively elicit NA responses because they present NA antigens in a large multivalent display with the proper topology and eliminate any potential loss from additional split vaccine manufacturing steps.

Our results showed that low amounts of inactivated virions readily elicit protective NA responses in mice and that the NA protective dose can be lowered by manipulating virions to increase NA content, confirming the importance of quantity. In addition to preventing severe weight loss, NA protected mice also developed strong antibody responses to challenge viruses with a heterotypic (different subtype) HA, supporting early reports on the benefits of the non-sterilizing immunity provided by NA [39]. Finally, we demonstrated that mixing inactivated virions with rNA resulted in stronger NA-based protection and higher NAI antibody responses than either component alone, more so for the homotypic (identical) than the heterotypic rNA. Together, these results demonstrate that inactivated virions are an adaptable vaccine platform with a high potential for eliciting protective antibody responses to both HA and NA.

Currently, many influenza vaccines are available from different antigen sources, however none have been combined into a single product. Our finding that BPL inactivated virions and insect cell produced recombinant NA can be used in a mix-and-shoot strategy suggests that at least some viral- and protein-based vaccines can be combined in a single syringe to improve protective responses to influenza antigens. While the benefits of combinatorial approach are clear for small molecule therapies, combining biologics remains challenging because of potential interference between antigens and excipient (buffer) incompatibility, either of which could potentially explain a previous observation that many available influenza vaccines blunted immune responses to a soluble rNA when they were combined [28].

In this study, we diluted inactivated virions and rNA in sterile PBS containing 1 mM CaCl_2_ to account for excipient compatibility and the two components were mixed briefly prior to injection due to our lack of stability data for the mixture. We also observed no evidence of antigen interference. Instead, mixing appeared to stimulate antibody responses to both homotypic and heterotypic rNAs, which were purified by two completely different approaches, and in one case the HA in the inactivated virion. For the homotypic rN1^BR18^ mixture there is likely an additive effect of the additional NA antigen, but the heterotypic rN2^KS17^ is presumably more reliant on the virion stimulation due to the lack of an inactivated H3N2 virion that would be present in any seasonal vaccine. However, we have not determined if the stimulation is due to the additional NA activity in the mixture or potential interactions between the two components. We also have not tested if the approach is compatible with other influenza or viral antigens, which can significantly expand potential applications of this mix and shoot strategy.

Coomassie stained gel densitometry revealed that NA comprises ∼10% of the total protein in the WSN^H1N1/BR18^ virions, which required ∼30 ng for protection, and ∼2.5% in the PR8^H1N1/BR18^ virions, which showed protection at the 125-ng dose. This implies that in our mouse model NA-based protection for N1 requires less than 5 ng of NA in a virion, whereas rN1 protection was only observed with 4 μg of protein alone or 1 μg combined with 30 ng of PR8^H1N1/BR18^ virions or poly I:C adjuvant. Based on this difference, full-length NA in a virion is a strikingly superior antigen at least compared to our soluble rN1, and if the properties of NA in a virion can be recapitulated it could dramatically improve any potential rNA vaccine.

Due to the potential benefits of NA, numerous studies have examined different NA vaccine approaches [45, 46]. However, most are proof-of-concept studies that used excessive antigen amounts in prime boost (2 dose) regimens, which can significantly mask antigen inferiority and often lack comparators. These studies also often provide little information about NA quality attributes (*e*.*g*. purity, activity, molecular structure) or effects of purification procedures or excipients on the structural conformation of NA, making reproducibility challenging. Despite these issues, many of these studies have been informative, but much like our prior work [27], they support a model where NA vaccines would be administered in addition to annual influenza vaccines, which is challenging from logistic or compliance perspectives. Certainly, our own data shows that adjuvanted rNA is potentially more protective than rNA mixed with inactivated virions, but the mixture is supplied in a single dose and has the added benefit of antibody responses against the HA in the virion. It is also easy to speculate that any stimulatory effect would likely increase in a quadrivalent influenza vaccine due to the higher virion amounts and adjuvants can be included in this formulation as well.

While our animal model produced reasonably consistent data, correlations between NAI titres and dose amounts are certain to change with the animal model, challenge virus dose, composition (*e*.*g*. sequences and subtypes) and NA viral amount, highlighting the difficulty of establishing a NA correlate of protection in humans. A good example is the protection results from the rN1 versus the rN2 immunization and challenge experiments where 4 μg of rN2 only provided partial protection when it was combined with virions, whereas 4 ug rN1 alone was fully protective. Our study also excluded any contributions from pre-existing immunity. Although tedious, these factors should be systematically investigated for individual NA subtypes if any additional efficacy is going to be achieved from including NA antigens in a vaccine.

One of the major reasons inactivated virions were replaced by split vaccines was the observed reactogenicity following administration in children. However, the elderly continues to be a major risk population for influenza where the potential reward from a more modern inactivated virion vaccine like the one recently tested [47] may confer efficacy that outweighs reactogenicity concerns. Clearly the benefits of combining rHA with inactivated virions or an optimal NA split vaccine formulation are easy to envision during a season where vaccines include a potentially detrimental egg adaptation in HA. They also should receive consideration during a potential pandemic due to the low immunoreactivity of some avian HA subtypes and their ability to be combined with other protective influenza antigens produced in different platforms.

## MATERIALS AND METHODS

### Reagents and viruses

Simple Blue Stain, Novex 4-12% Tris-Glycine SDS-PAGE gels, Immulon 2HB flat bottom 96-well microtiter plates, Maxisorp 96-well plates, Sf-900™ II SFM, and 1M Dithiothreitol (DTT) were obtained from Thermo Fisher Scientific. Ultra-clear ultracentrifuge tubes were purchased from Beckman Coulter. Specific-pathogen-free (SPF) eggs and DBA/2J mice (#800671) were purchased from Charles River Labs and Jackson Labs (Bar Harbor, ME), respectively. Bovine fetuin, bovine serum albumin (BSA), Tween 20, *o*-phenylenediamine dihydrochloride (OPD), beta-propiolactone (BPL), HRP-linked peanut agglutinin, phosphate-citrate buffer with sodium perborate and Methyl-α-D mannose were purchased from Sigma. Lentil lectin resin, SP HP resin, HiTrap Q HP Anion Exchange Columns, HisTrap FF columns, and Vivaspin 2 Molecular weight cut-off (MWCO) 30 kD protein concentrators were purchased from Cytiva (Marlborough, MA). Turkey red blood cells (TRBCs) and 2’-(4-methylumbelliferyl)-α-d-*N*-acetylneuraminic acid (MUNANA) were acquired from Poultry Diagnostic and Research Centre (Athens, GA) and Cayman Chemical, respectively. Dulbecco’s phosphate buffered saline (DPBS, pH 7.4) and Phosphate buffered saline (PBS, pH 7.2 and 7.4) were obtained from KD Medical. KPL Coating Solution Concentrate (10X) was purchased from Seracare Life Sciences Inc. and poly(I:C) (Polyinosinic-polycytidylic acid) from InvivoGen. Descriptions of the IAVs used in this study are listed in Table 1, which include the A/Brisbane/02/2018 (H1N1) strain (BR18^H1N1^) kindly provided by WHO Collaborating Center in Melbourne Australia.

**Table 1.**
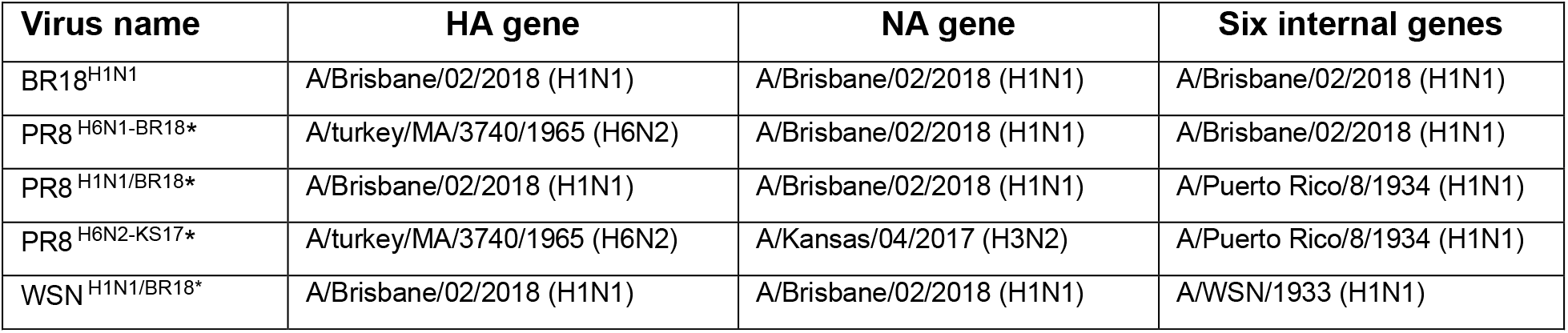
Strain origin of influenza genes in the viruses used for this study. All viruses were propagated in 10-day-old SPF eggs. ^*^ Viruses were generated by reverse genetics as previously described [38].

### Virus propagation, BPL inactivation and isolation

Viruses were propagated by inoculating 10-day old embryonated SPF eggs with 100 μl of a 1/1000 virus dilution in PBS pH 7.4. Infected eggs were incubated at 33°C for 3 days and placed at 4 °C for 2 h prior to harvesting and clarifying the alantoic fluid by centrifugation (2000 × g; 5 min) at 4 °C. Challenge viruses (PR8^H6N1-BR18^, PR8^H6N2-KS17^ and BR18^H1N1^) in clarified alantoic fluid were directly aliquoted and stored at -80 °C. PR8^H1N1/BR18^ and WSN^H1N1/BR18^ viruses in alantoic fluid were inactivated with 0.1% BPL overnight at 4 °C and transferred to ultracentrifuge tubes, under layered with a 4 ml cushion (25% sucrose (w/v), PBS pH 7.4 and 1 mM CaCl_2_) and sedimented (100,000 × g; 45 min) in a SW41Ti swinging bucket rotor at 4 °C. Sedimented virions were re-suspended in PBS pH 7.2 containing 1 mM CaCl_2_ and protein concentrations were determined using a BCA protein assay kit (Pierce). Resuspended viruses were adjusted to 1 mg/ml with PBS pH 7.2 containing 1 mM CaCl_2_ aliquoted (100 μl) and stored at -80 °C. Inactivation of virions used for immunization was confirmed by the absence of HA units (HAU) titres and NA activity in the alantoic fluid of eggs after three consecutive passages at a 1:10 dilution.

### Sodium dodecyl-sulfate polyacrylamide gel electrophoresis and Coomassie staining

Isolated viruses (∼5 μg total viral protein) were mixed with Tris-Glycine SDS sample buffer (2X) containing 0.1 M DTT as indicated, heated at 50 °C for 5 min and resolved on 4-12 % polyacrylamide Tris-Glycine SDS-PAGE wedge gels at 150V for 1 h. Gels were washed three times for 5 min with dH_2_O, stained with simple blue for 1 h and destained with dH_2_O prior to imaging with an Azure C600 Bioanalytical Imaging System (Azure Biosystems).

### Recombinant N1 (rN1) and N2 (rN2) constructs

All recombinant NAs were produced by infecting insect cells with recombinant baculoviruses (BVs) that encode for secreted rN1 or rN2 variants. rN1 used for immunization was encoded by a signal peptide (GP67a), 6×His-tag, tetrabrachion tetramerization domain, followed by a 7-residue linker and N1 residues 35–469 from the IAV strain A/Brisbane/02/2018 (H1N1). rN1 used for ELLA was identical except for the tetramerization domain which was from the human vasodilator stimulating protein (VASP). rN2 was encoded by a signal peptide (azurocydin), followed by a 13-residue tag (Spy), tetrabrachion tetramerization domain, 7-residue linker and N2 residues 35–469 from the IAV strain A/ Kansas/14/2017 (H3N2).

### Purification of rN1 and rN2

Sf9 insect cells grown at 27.5°C to a density of 2-4 × 10^6^ cells/ml in shaker flasks at 120 rpm were infected with BVs (2% v/v) for 72 – 96 h prior to clarifying the culture supernatant by centrifugation (10,000 × g; 30 min). rN1 used for immunization was purified from the supernatant with a HisTrap FF 1-ml column as previously described [27]. rN2 was purified by multistep chromatography. The supernatant was adjusted to pH 7.5 with NaOH, loaded onto a 10 ml Lentil lectin agarose column using an AKTA start and washed with 15 column volumes (CVs) of buffer A (30 mM Tris pH 7.5, 1.0 mM CaCl_2_, and 0.5 M NaCl) before eluting bound proteins with buffer A containing 0.5 M Methyl-α-D mannose (Sigma). Fractions containing NA activity were pooled, 60 volumes of buffer B (30 mM Tris pH 7.5 containing 1.0 mM CaCl_2_) were added and the sample was concentrated with a Vivaspin 2 MWCO 30 kD concentrator prior to loading onto a HiTrap Q HP 1-ml column and washing with 15 CVs buffer B containing 50 mM NaCl. rN2 was eluted in 20 CVs buffer B containing 200 mM NaCl, dialyzed overnight at 4 °C against 50 volumes buffer C (30 mM MES pH 6.5 containing 1.0 mM CaCl_2_). The Q elution was loaded onto a 1-ml SP HP resin column, washed with 15 CVs buffer C containing 50 mM NaCl and the rN2 was eluted in 20 CVs buffer C containing 200 mM NaCl. The rN2 was dialyzed overnight at 4 °C against 50 volumes storage buffer (30 mM Tris pH 6.5, 1.0 mM CaCl_2_, 150 mM NaCl, and 2.0 % Glycerol), aliquoted and frozen at -80 °C. rN1 used for ELLA was purified using Lentil lectin agarose and a HisTrap FF 1-ml column followed by dialysis against storage buffer and storage at -80 °C.

### Characterization of rN1 and rN2

rNA protein concentrations were determined by Absorbance (Abs) at 280 nm using the molar extinction coefficients for rN1 (105850 M^-1^cm^-1^) and rN2 (83975 M^-1^cm^-1^), respectively. Purity was assessed by Coomassie staining following SDS-PAGE. NA activity was measured using MUNANA as previously described [48] and the oligomeric state was monitored by size exclusion chromatography using an Agilent 1260 prime HPLC equipped with an AdvanceBio SEC 300Å column and a variable wavelength detector set at 220 and 280 nm with PBS pH 7.2 containing 1 mM CaCl_2_ as the mobile phase and a flow rate of 1 ml/min. AdvanceBio SEC 300Å protein standard (Agilent) was included to correlate the elution volume with molecular weight.

### Ethics statement

All animal experiments were approved by the U.S. FDA Institutional Animal Care and Use Committee (IACUC) under Protocol #2003–18. The animal care and use protocol meets National Institutes of Health (NIH) guidelines.

### Median lethal dose (LD_50_) determination

Female DBA/2J mice were housed in specific-pathogen-free conditions. Nine-week-old mice (5 per group) were intranasally infected with ten-fold serial dilutions of BR18^H1N1^, PR8^H6N1-BR18^ or PR8^H6N2-KS17^ viruses in PBS pH 7.2 and mice were monitored for clinical and weight changes for 2 weeks post-infection. The LD_50_ was determined based on the survival (<25% weight loss) ratio at each dose. Sera was collected from mice that received the highest virus dose without exhibiting 25% weight loss to evaluate NA and HA antibody responses following infection. Mice were euthanized with CO_2_ at a rate of at least 30% per minute following the institutional Animal Study Proposal (ASP).

### Intramuscular (I/M) immunizations with inactivated virions and recombinant NA

Mice were earmarked prior to immunization to correlate serological and viral challenge results to specific mice. BPL inactivated PR8^H1N1/BR18^ and WSN^H1N1/BR18^ virions were thawed and diluted in sterile PBS pH 7.2 containing 1 mM CaCl_2_ to protein concentrations that provided the indicated dose amounts in 50 μl. Immunizations were performed by I/M administration of the 50 μl dose to six-week-old DBA/2J female mice (5 or 3 per group) with a 0.5 ml U-100 syringe equipped with a 28.5 gauge needle (Exelint International Co.). For experiments involving recombinant N1 (rN1) or N2 (rN2) the indicated protein amounts were either brought up directly to 50 μl with sterile PBS pH 7.2 containing 1 mM CaCl_2_ or combined with 0.03 μg of BPL inactivated PR8^H1N1/BR18^ virions, or 1 μg of poly(I:C), and then brought up to 50 μl with sterile PBS pH 7.2 containing 1 mM CaCl_2_. Sterile PBS pH 7.2 with 1 mM CaCl_2_ was used as a control in all immunization experiments. Sera were obtained from the mice at two and/or three weeks post-immunization for serological analysis.

### Viral challenge

After sera collection three weeks post-immunization, mice were intranasally infected with 10 ×LD_50_ of PR8^H6N1-BR18^, PR8^H6N2-KS17^ or BR18^H1N1^ virus diluted in 50 μl of sterile PBS pH 7.2. Changes in weight were recorded for two weeks post-infection and moribund mice or mice that showed 25% weight loss were euthanized by CO_2_ in accordance with the ASP. Sera was collected from surviving mice by cardiac bleed 2 weeks post-challenge.

### NA inhibitory antibody (NAI) titre determined by fetuin-based enzyme-linked lectin assay (ELLA)

ELLA was used to determine NAI titres in the collected sera samples as previously described [23]. Briefly, serum samples were diluted two-fold in 96-well plates using DPBS pH 7.4 containing 1% BSA and 0.5% Tween 20, and 50 μl from each well was transferred to a 96-well fetuin-coated (2.5 μg/well) Maxisorp plate. Each well than received 50 μl of a predetermined dilution of PR8^H6N1-BR18^ virus or rN1 protein in DPBS pH 7.4 containing 1% BSA and 0.5% Tween 20 that produce an Abs 490 nm of ∼2.0 in the absence of sera. Plates were incubated overnight at 37°C, washed six times with PBS pH 7.4 containing 0.05% Tween 20 (PBST) and incubated 2 h at room temperature with 100 μl of HRP-conjugated peanut agglutinin diluted to 1 μg/ml in DPBS pH 7.4 containing 1% BSA. Plates were washed with PBST, 100 μl of 0.5 mg/ml OPD in 50 mM phosphate citrate pH 5.0 buffer was added and wells were developed 10 min at room temperature. Reactions were stopped with 100 μl of 1 N H_2_SO_4_ and the Abs 490 nm was read using a Biotek Cytation 5. Wells that received only buffer (no sera, virus or protein) were averaged to determine the background. Background corrected Abs 490 nm values were plotted with respect to the reciprocal sera dilution in GraphPad Prism 8 software and curves were generated using the standard curve, sigmoidal, 4 parameter logistic function with least squares fit to determine the NAI titer which corresponds to the 50% inhibitory concentration or IC_50_.

### NA antibody (NA) titre determined by enzyme-linked immunosorbent assay (ELISA)

Purified PR8^H6N2-KS17^ virions (1 mg/ml) were diluted 1:200 in 1×Coating Buffer, 100 μl was transferred to each well of 96-well 2HB ELISA plate (0.5 μg/well) and the plate was incubated at 4 °C overnight. The coating solution was removed, 200 μl of blocking solution (1% BSA) was added to each well and the plate was incubated at 37 °C for 1 h. Blocking solution was removed and 100 μl of prepared two-fold sera dilutions in 0.1% BSA-PBST was added to each well. Plates were incubated at 37 °C for 1 h, wells were washed 3 times with 200 μl PBST and 100μl of Anti-mouse IgG-HRP secondary antibody diluted 1:20,000 in 0.1% BSA-PBST was added to each well. Plates were incubated 1 h at 37 °C, wells were washed 3 times with PBST and developed for 10 min at room temperature after adding 100 μl of 0.5 mg/ml OPD diluted in 50 mM phosphate citrate pH 5.0 buffer. Reactions were stopped by adding 100 μl of 1 N H_2_SO_4_ and the Abs 490 nm was read using a Biotek Cytation 5. ELISA titres were reported as the lowest sera dilution with an Abs 490 nm value two times background (∼0.1).

### Hemagglutinin Inhibition assay (HAI)

Each sera sample (10 μl) was treated with 30 μl of Receptor Destroying Enzyme (RDE) Kit (Cosmos Biomedical Ltd.) overnight at 37 °C and heat inactivated at 56 °C for 45 min. Treated sera was brought up to 100 μl using PBS pH 7.2. Serial two-fold dilutions of the 1/10 treated sera were prepared in 96-well plates containing 25 μl of PBS pH 7.2 in each well prior to adding 25 μl of the indicated viruses (BR18^H1N1^ or PR8^H6N1-BR18^) diluted in PBS pH 7.2 to 4 HAU, which was predetermined. Plates were incubated at room temperature 30 min prior to adding 50 μl of 0.5% TRBCs in PBS pH 7.2 and an additional 45 min incubation. HAI titres were calculated based on the lowest reciprocal sera dilution that prevented hemagglutination.

### Statistical analysis

Statistical analysis was performed with GraphPad Prism 8 software with the assumptions that the data follows a Gaussian distribution. Student’s unpaired t-test was performed using a two-tailed analysis and a confidence interval (CI) of 95%.

## DATA AVAILABAILITY

All relevant data are within this manuscript and its Supporting Information files.

## ADDITIONAL INFORMATION

**Supplementary information -** The supplementary data includes two figures and a supplementary data file with all of the raw data presented in the manuscript.

**Reprints and permission information**.

**Correspondence and requests for material -** should be addressed to R.D.

## ACKNOWLEDGEMENTS

We would like to thank members of the Division of Viral Products at the Center for Biologics Evaluation and Research (CBER), especially Dr. Hana Golding and Dr. Amy Rosenfeld for the critical reading of the manuscript and offering several helpful suggestions. We are grateful to the entire staff of the Division of Veterinary Services at CBER for extraordinary animal care, especially Amy Guardado for technical support and advice. We also thank Dr. Luca Giurgea (NIH) and Dr. Matthew Memoli (NIH) for useful discussions on the implications of the data. This work was supported by Intramural funding to R.D. from CBER at the US Food and Drug Administration.

## AUTHOR CONTRIBUTIONS

All author contributions are an informal communication and represent their own best judgment. These comments do not bind or obligate FDA. R.D. conceived and supervised the study. M.R.M. performed the experiments and analyzed the data. J.G. provided all the reassortant viruses and assisted with HAI assays. H.W. provided rN1^BR18^ and assisted with ELLA assays. H.K. provided rN2^KS17^ and helped with the rN1 and rN2 analysis. R.D., M.R.M. and L.K. wrote and edited the manuscript with input from all authors.

---

**Supplemental Figure 1.**
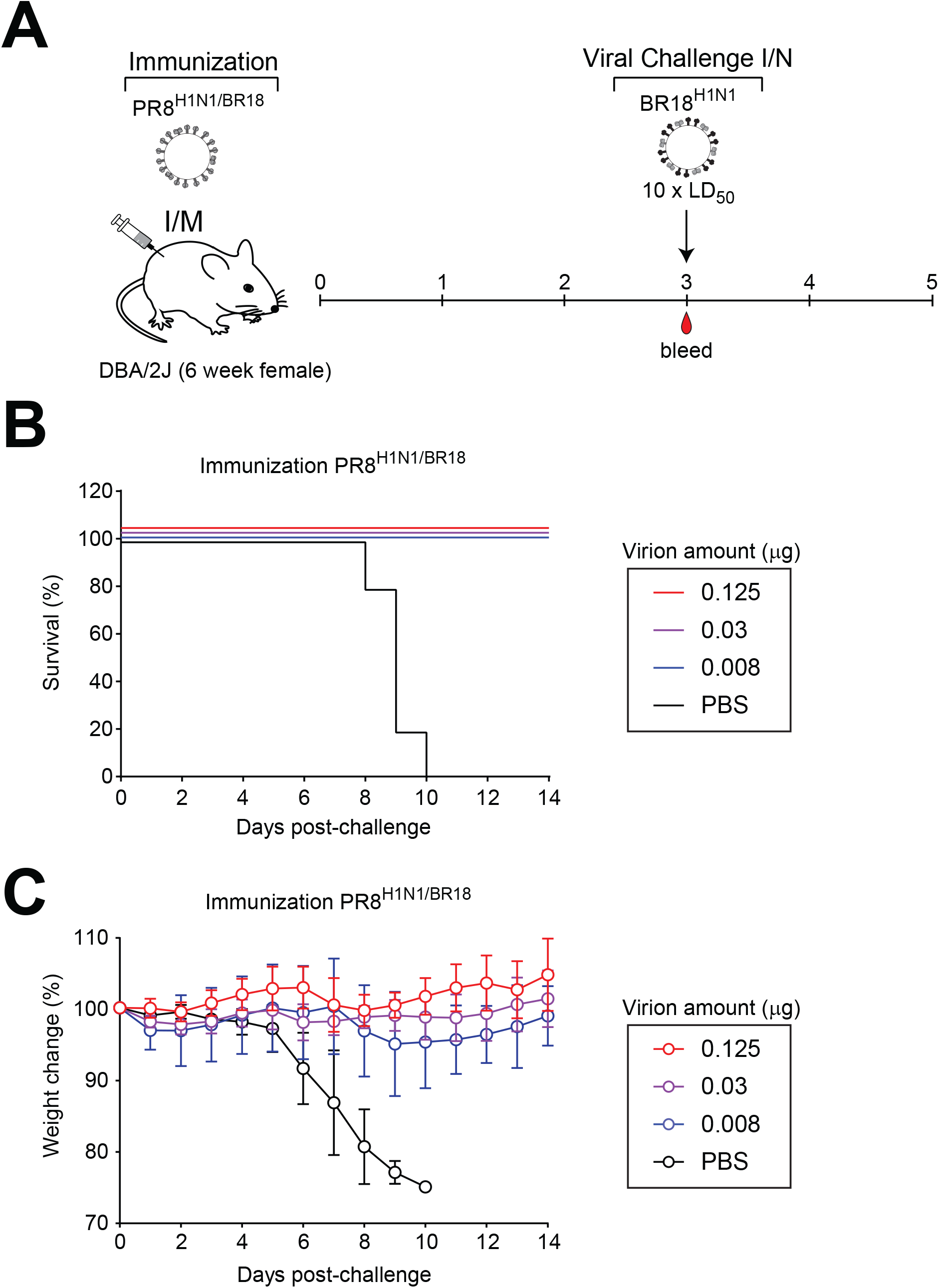
Immunization and challenge with virions containing identical H1 and N1 antigens from BR18. **A**. Diagram of the lethal viral challenge for evaluating the protection provided by BPL inactivated PR8^H1N1/BR18^ virions against a virus (BR18^H1N1^) carrying identical H1 and N1 antigens. **B**. Graph displaying the percentage of mice (n=5) in each immunization group that survived the BR18^H1N1^ virus challenge. **C**. Body weight was monitored in all mice following the viral challenge and the mean for each vaccine group is displayed ± s.d.

**Supplemental Figure 2.**
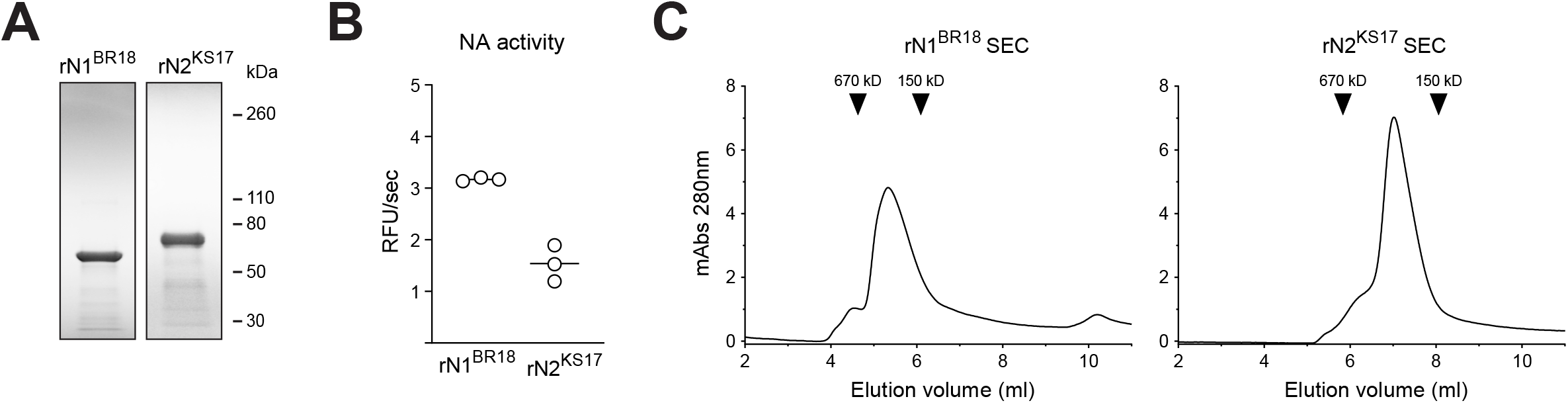
Characterization of the rN1^BR18^ and rN2^KS17^ antigens. **A**. Image of a reducing Coomassie stained SDS-PAGE gel (12-4%) containing 4 μg of the rN1^BR18^ and rN2^KS17^ antigens. **B**. Enzymatic activity of the rN1^BR18^ and rN2^KS17^ antigens (100 ng/well) is displayed as relative fluorescent units (RFU) per second. Activity was measured in 200 μl containing 50 μM MUNANA and calculated by the slope of the curve during a 10 min reaction. **C**. Size exclusion chromatography (SEC) profiles of the rN1^BR18^ and rN2^KS17^ antigens monitored by absorbance (Abs) at 280 nm. Elution peaks for the indicated molecular weight standards are depicted by arrowheads.

